# An In Vivo Method for Diversifying the Functions of Therapeutic Antibodies

**DOI:** 10.1101/2020.12.12.421834

**Authors:** Ming Tian, Hwei-Ling Cheng, Michael T. Kimble, Kelly McGovern, Peyton Waddicor, Yiwei Chen, Elizabeth Cantor, Mengting Qiu, Marie-Elen Tuchel, Mai Dao, Frederick W. Alt

## Abstract

V(D)J recombination generates mature B cells that express huge repertoires of primary antibodies as diverse immunoglobulin heavy (IgH) and light chains (IgL) of their B cell antigen receptors (BCRs). Cognate antigen binding to BCR variable region domains activates B cells into the germinal center (GC) reaction in which somatic hypermutation (SHM) modifies primary variable region-encoding sequences, with subsequent selection for mutations that improve antigen-binding affinity, ultimately leading to antibody affinity maturation. Based on these principles, we developed a humanized mouse model approach to diversify an anti-PD1 therapeutic antibody and allow isolation of variants with novel properties. In this approach, component Ig gene segments of the anti-PD1 antibody underwent *de novo* V(D)J recombination to diversify the anti-PD1 antibody in the primary antibody repertoire in the mouse models. Immunization of these mouse models further modifies the anti-PD1 antibodies through SHM. Known anti-PD1 antibodies block interaction of PD1 with its ligands to alleviate PD1-mediated T cell suppression, thereby boosting anti-tumor T cell responses. By diversifying one such anti-PD1 antibody, we derived many new anti-PD1 antibodies, including anti-PD1 antibodies with the opposite activity of enhancing PD1/ligand interaction. Such antibodies theoretically might suppress deleterious T cell activities in autoimmune diseases. The approach we describe should be generally applicable for diversifying other therapeutic antibodies.

**Significance Statement:** The immune system is capable of producing enormous varieties of antibodies to counter pathogenic infections. In this study, we devised a method to harness the power of the immune system to produce potentially therapeutic antibodies. As a test, we used this method to generate antibodies against the human PD1 molecule, a negative regulator of T cell activity. Available anti-PD1 therapeutic antibodies inhibit the function of PD1, thereby boosting T cell activity against tumor cells. Through our approach, we diversified one such anti-PD1 antibody in our mouse models. From this work, we obtained multiple new anti-PD1 antibodies, some of which can stimulate, rather than inhibit, PD1 function *in vitro*. Such PD1 stimulatory antibodies could potentially be used to suppress pathogenic T cell activities in autoimmune diseases. This method could be used to diversify the functions of other therapeutic antibodies.

## Introduction

Therapeutic antibodies must meet stringent criteria for clinical application (1, 2). For this reason, lead antibodies generated by various antibody development platforms often may benefit from further modifications. Toward this end, we developed an *in vivo* method for antibody diversification and optimization. Our approach exploits antibody diversification mechanisms during B cell development and activation in mice (3). At the progenitor B cell stage, V(D)J recombination joins Ig VH, D and JH gene segments into complete exons that encode IgH variable regions of antibodies and similarly joins the VL and JL segments that encode the variable regions of IgL chains of antibodies. A major portion of antibody diversity comes from mechanisms that diversify the junctions of V, D, and J segments during V(D)J recombination. Thus, as the portions of the V_H_-D and D-J_H_ junctions of antibody IgH chain variable regions or the V_L_-J_L_ junctions of IgL chain are part of the antigen-contact complementarity-determining region 3 (CDR3) of IgH and IgL chains, junctional diversification generates enormous varieties of primary antigen-binding B cell receptors (BCRs) for any given combination of germline encoded V, D and J segments (3). Unique BCRs are displayed on the surface of each clonally generated primary B cell, which then migrate to peripheral lymphoid tissues. There, antigen binding to a cognate B cell receptor stimulates the corresponding B cells, which ultimately can participate in germinal center (GC) reactions (4). The Ig variable regions of GC B cells accumulate SHMs that can further diversify IgH and IgL CDR3 sequences, as well as the two other antigen-contact CDR1 and CDR2 encoded in germline V_H_ and V_L_ gene segments (4). Some SHMs increase antigen-binding affinity of the B cell receptor, and the germinal center microenvironment selects for B cells with increased antigen-binding affinity. Repeated cycles of mutation and selection can lead to antibody affinity maturation (4).

For our new approach, we generate mice that predominantly assemble the IgH V(D)J exon of an existing therapeutic antibody by V(D)J recombination during B cells development, creating vast repertoires of primary B cells expressing different variations of the antibody due to junctional diversification of the IgH variable region CDR3. Then we immunize these mice with the cognate antigen for the therapeutic antibody to further diversify the primary antibody sequences by SHM and affinity maturation in the GC. Relative to *in vitro* antibody development platforms, such as phage display (5–7) or yeast display (8), we hypothesized that our *in vivo* approach could yield some antibodies that are more suitable for clinical applications. For example, B cell developmental checkpoints can eliminate poly-reactive antibodies (9). Moreover, B cell survival depends on functional B cell receptors (10); this requirement selects *in vivo* for antibodies with stable and normal conformations. For clinical application, antibodies are usually produced in mammalian cells. When antibodies are expressed in mouse B cells, functional properties selected in mice may be more likely to be preserved than those for *in vitro* evolved antibodies from bacteria or yeast (5–8).

As a proof of principle, we tested the ability of our *in vivo* antibody diversification approach to generate new versions of anti-PD1 antibody from an existing prototype (11). PD1 is a surface receptor on activated T cells (12). Interaction of PD1 with its ligands represses T cell activity (13, 14). By blocking the association between PD1 and its ligands, antibodies against PD1 or its ligand can neutralize the PD1-dependent negative regulatory pathway, thereby boosting T cell activity (15–18). Based on this function, anti-PD1 antibodies have proved effective in cancer immunotherapy (19–23). It is conceivable that further diversification of existing anti-PD1 antibodies could potentiate their efficacy for current uses or alter their activity for new applications. Based on the *in vivo* diversification approach, we have generated a set of new anti-PD1 antibodies, several of which actually enhance, rather than block, PD1 interaction with its ligand.

## Results and Discussion

### Generation of mouse models to diversify an anti-PD1 antibody

We chose to diversify an anti-PD1 antibody 17D8 (11) to produce variant antibodies with new properties. The 17D8 anti-PD1 antibody was developed by Bristol-Meyers Scripps (11) and is homologous to the Nivolumab cancer immunotherapy antibody (19, 21). Besides its obvious clinical significance, the technical rationale for choosing the 17D8 antibody is that our approach could generate 17D8-related antibodies with an enormous diversity of IgH variable region CDR3s (“CDR H3”), which could alter CDR H3-encoded antigen recognition properties. In addition, since the IgH or IgL variable region sequences of the 17D8 antibody are close to the germline versions of the component Ig gene segments (11), the antigen-recognition specificity and affinity of this antibody might be further modified by SHM. To rapidly generate a mouse model system to test our 17D8 in vivo antibody diversification approach, we introduced all necessary genetic modifications (see below) into mouse embryonic stem (ES) cells and then injected the engineered ES cells into Recombination Activating Gene-2 (RAG-2) deficient blastocysts to generate chimeric mice, in which all mature lymphocytes develop from the injected ES cells with the entire set of *in vitro*-introduced genetic modifications (24). With this RAG-2 deficient blastocyst complementation (RDBC) approach, we can use the chimeric mice directly for immunization experiments, thereby saving the expense and time associated with conventional germline breeding (25).

To diversify the 17D8 antibody via junctional diversification during progenitor B cell development, we separated its IgH variable exon into its component VH and recombined DJ_H_ segments, derived from human V_H_3-33, D1-1 and J_H_4 segments, respectively. We then substituted the 17D8 V_H_ segment for the most proximal mouse V_H_81X gene segment in ES cells, in which we had deleted the IGCRI regulatory region to render the integrated human 17D8 V_H_ segment by far the most utilized V_H_ in developing B cells of RDBC chimeras derived from these ES cells (Figure 1A) (25, 26). To further enrich the representation of the 17D8 V_H_ and DJ_H_ segments in primary antibody repertoire, we also deleted the J_H_ region of the other IgH allele in the targeted ES cells (Figure 1A). To reconstitute the complete 17D8 HC variable region during V(D)J recombination in developing RDBC progenitor B cells, we replaced the mouse DQ52-J_H_ region in the modified ES cells with the recombined 17D8 DJ_H_ segment. A recombination signal sequence (RSS), upstream of the 17D8 DJ_H_ segment, enables the joining of the 17D8 DJ_H_ segment with upstream V_H_ segments, including the 17D8 V_H_ segment in developing mouse progenitor B cells (Figure 1A). Because the RSS of the assembled 17D8 DJ_H_ segment was derived from a mouse D segment, the 12/23 rule of V(D)J recombination (27) restricts the recombination of the 17D8 DJ_H_ segment RSS to upstream V_H_ RSS, but is not the incompatible D RSS for joining.

**Figure 1.**
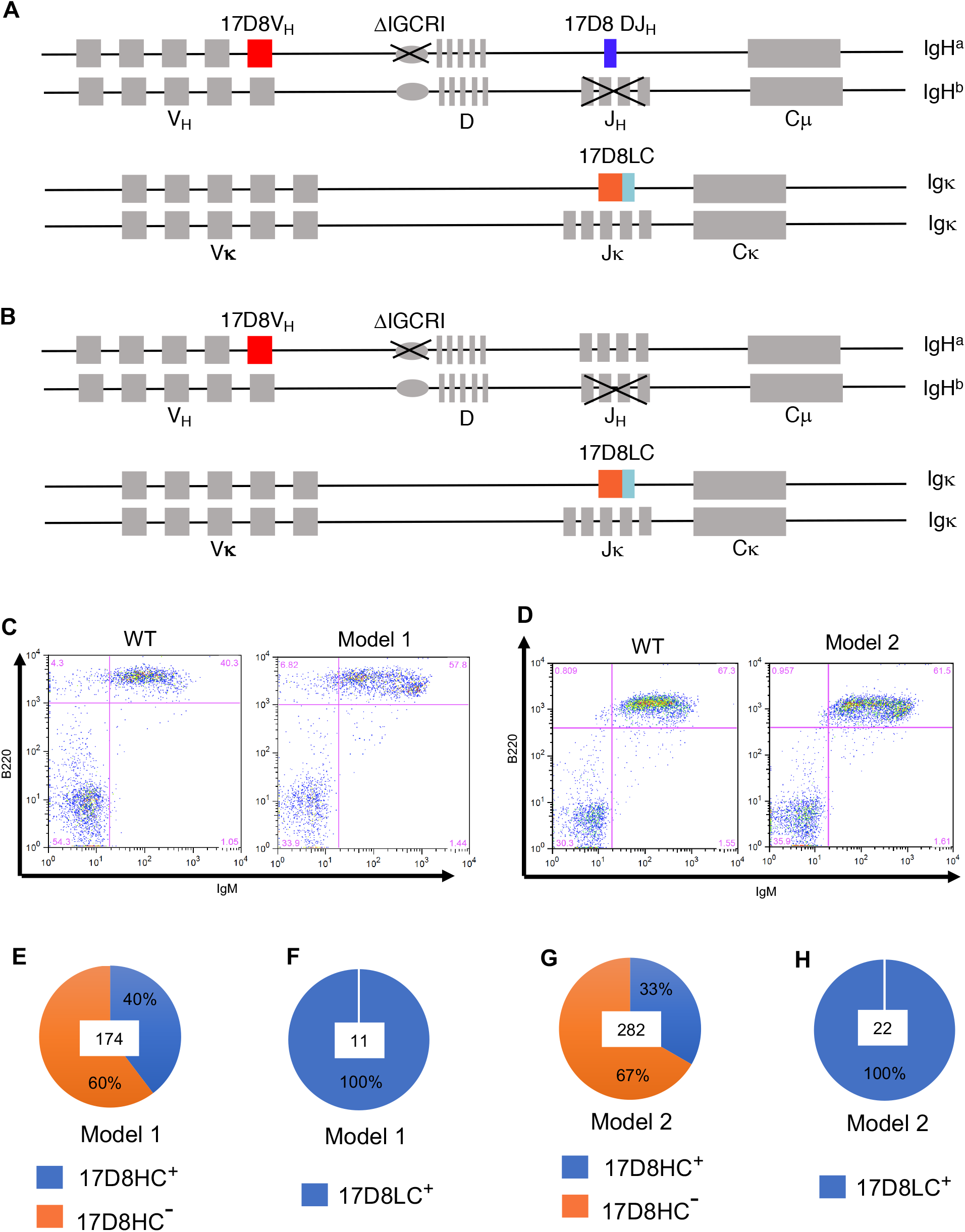
Description of the mouse models to diversify an anti-PD1 antibody, 17D8. **A.** Diagram of mouse model 1. See text for details. **B.** Diagram of mouse model 2. See text for details. **C.** FACS analysis of B cells in blood from wild-type 129/Sv mouse and mouse model 1. The surface markers detected by the FACS staining are indicated next to the axis. **D.** FACS analysis of B cells in blood from wild-type 129/Sv mouse and mouse model 2. The surface markers detected by the FACS staining are indicated next to the axis. **E.**The percentage of IgH rearrangements involving the 17D8 V_H_ and DJ_H_ segments in mouse model 1. Hybridomas were generated from splenic B cells of mouse model 1. IgH rearrangements involving the 17D8 V_H_ and DJ_H_ segments in each hybridoma clone were detected with PCR. In total, 174 hybridoma clones, the number at the center of pie chart, were analyzed, and 40% of the clones were positive for 17D8 V_H_(D)J_H_ recombination products (17D8HC^+^). **F.** The fraction of Igκ transcripts corresponding to 17D8LC in mouse model 1. 5’RACE was performed from Cκ, using RNA from splenocytes of mouse model 1. 11 PCR products, the number shown at the center of the pie chart, were sequenced, and all of them corresponded to the 17D8LC. **G.** The percentage of IgH rearrangements involving the 17D8 V_H_, mouse D and J_H_ segments in mouse model 2. IgH rearrangements involving the 17D8 V_H_, mouse D and J_H_ segments in each hybridoma clone were detected with PCR. In total, 282 hybridoma clones, the number at the center of pie chart, were analyzed, and 33% of the clones contained rearrangements involving the 17D8 V_H_. **H.** The fraction of Igκ transcripts corresponding to 17D8LC in mouse model 2. 5’RACE was performed from Cκ, using RNA from splenocytes of mouse model 1. 22 PCR products, the number shown at the center of the pie chart, were sequenced, and all of them corresponded to the 17D8LC.

After V(D)J recombination, the V_H_-DJ_H_ junction can gain or lose random nucleotides; such junctional diversification can create tremendous variations in CDR H3 (28). Thus, V(D)J recombination of 17D8 V_H_ with DJ_H_ *in vivo* does not simply regenerate the original 17D8 antibody heavy chain (HC); the process will produce a large library of related HCs that differ in CDR H3 length and/or sequence. Some of the new CDR H3s may recognize PD1 with equal or higher affinity than the original CDR H3 in 17D8 or even target different epitopes. We refer to mice generated with this diversification strategy as PD1 Diversification Mouse Model 1. In a variation of this strategy, referred to as PD1 Diversification Mouse Model 2, we did not incorporate the 17D8 DJ_H_ segment into the mouse IgH locus (Figure 1B), allowing the 17D8 V_H_ segment to undergo V(D)J recombination with mouse D and J_H_ segments, with junctional diversification occurring at both V_H_-D and D-J_H_ joints. As a result, the CDR H3 diversity in this model should be even higher than that in PD1 Diversification Mouse Model 1 with the 17D8 pre-rearranged DJ_H_ segment. A potential caveat of the PD1 Diversification Mouse Model 2 strategy is that antibodies generated in this model will contain mouse D and J_H_ segments in their CDR H3s, which in theory might provoke immune response in humans. Mitigating this concern, CDR H3 often retains minimal remnants of the D segment, and mouse J_H_s are homologous to human counterparts. Furthermore, due to junctional diversification, human antibodies also contain highly heterogenous CDR H3s. So, mouse CDR H3s may not pose a major problem for human applications.

To complement 17D8 HC expression, we integrated a pre-rearranged (VκJκ) variable region (V) exon for the 17D8 light chain (LC) into the mouse Jκ locus (Figure 1A and 1B) in ES cells, which already harbored the modified IgH locus for PD1 Diversification Mouse Model 1 or 2. The resultant ES clones, now engineered to express both diversified 17D8 HCs and a CDR L3-prefixed 17D8 LC were used to generate chimeric mice by RDBC (24). CDR L3 is much less diverse than CDR H3, because CDR L3 lacks a D segment and involves only V_L_-J_L_ junctions. In addition, CDR H3s are tremendously diversified via terminal deoxynucleotide transferase (TdT)-mediated N region additions to recombination junctions (29); while CDR L3s in mouse IgL variable regions are not diversified by N-nucleotide addition, due to the absence of TdT expression in mouse precursor-B cells that undergo V_L_ to J_L_ joining (30). For these reasons, we did not aim to diversify the 17D8 CDR L3 through V(D)J recombination in this set of mouse models. In addition, one advantage of expressing a pre-rearranged human V exon for the 17D8 LC is that, due to negative feedback regulation from the expressed rearranged knock-in IgL chain (31, 32), most naïve B cells in the mouse model should express the 17D8 LC. Thus, in conjunction with the dominance of the diverse 17D8-derived HCs, the majority of B cells in these two mouse models should express 17D8-related antibodies with a diverse range of CDR H3s.

Based on FACS analysis of blood lymphocytes, the RDBC chimeric mice for both mouse models had comparable B cell populations as wild-type 129/Sv mice (Figure 1C and 1D). As explained above, a large fraction of B cells should express HCs containing the 17D8 V_H_ and DJ_H_ segments in PD1 Diversification Mouse Model 1, or 17D8 V_H_ segment in association with mouse D and J_H_ segments in PD1 Diversification Mouse Model 2. To confirm this expectation, we generated hybridomas from splenic B cells of these mice. Indeed, 40% of B cell hybridomas from PD1 Diversification Mouse Model 1 contained 17D8 V_H_-DJ_H_ recombination products (Figure 1E). Since the J_H_ region has been deleted from the other IgH allele, all the 17D8 HC rearrangements should be productive and support B cell survival. Similarly, in PD1 Diversification Mouse Model 2, the 17D8 V_H_ segment recombined with mouse Ds and J_H_s in approximately 33% of B cell hybridomas (Figure 1G). Due to feedback regulation from the knock-in 17D8 variable exon-encoded IgL chain, rearrangement of the endogenous Igκ locus was inhibited in developing B cells in the RDBC chimeric mice, and all the Igκ transcripts from the splenic B cells of both mouse models corresponded to the 17D8 LC (Figure 1F and 1H).

### Isolation of new anti-PD1 antibodies from the mouse models

In both PD1 Diversification Mouse Models, V(D)J recombination diversifies the CDR H3 of the 17D8 antibody. At least a subset of the new CDR H3s may be able to interact with PD1, but with different affinities or targeting distinct epitopes from the original CDR H3 in the 17D8 antibody. Immunization of these mouse models with PD1 could activate naïve B cells that express new anti-PD1 antibodies. Some of the activated B cells will undergo SHM (33) that might further influence their PD1 binding affinity. For immunization of these mouse models, we used a fusion protein that consists of the extracellular domain of the human PD1 and Glutathione-S-transferase (GST). The GST portion facilitates affinity purification of the protein and may provide extra epitopes for helper T cells, which are critical for affinity maturation during GC reaction (34). In this regard, thymic tolerance mechanisms should purge T cells that recognize mouse PD1 epitopes, as well as homologous epitopes in human PD1. The exogenous GST could compensate for the potential paucity of helper T cells (34) that recognize antigenic peptides from the human PD1. Immunization with PD1-GST could induce antibodies against both the PD1 and GST portion of the fusion protein. To specifically detect anti-PD1 antibodies in ELISA, we used PD1 fused to human IgG4 fragment crystallizable (Fc) region; this fusion protein lacks the GST portion of the immunogen and should only interact with anti-PD1 antibodies from the immunized mice. Serum from unimmunized mice did not contain detectable PD1 binding activity in ELISA (Figure 2A and 2B). Two rounds of immunization with PD1-GST induced robust anti-PD1-binding IgG in plasma from both mouse models (Figure 2A and 2B).

**Figure 2.**
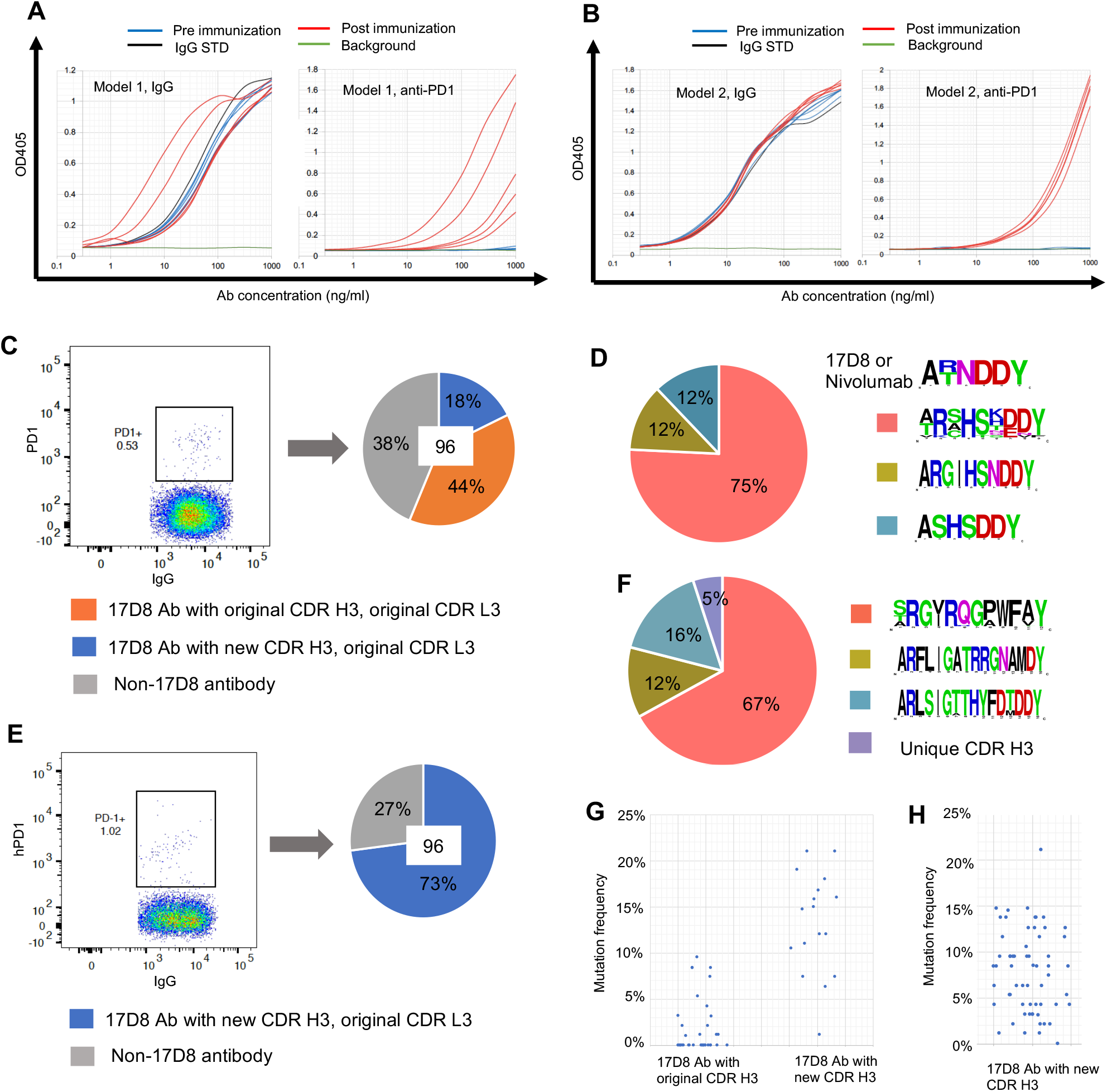
Immunization and isolation of new anti-PD1 antibodies from mouse model 1 and 2. **A.** ELISA detection of anti-PD1 IgG in plasma after immunization of mouse model 1. The plot on the left shows IgG concentrations in mouse plasma, collected two weeks after the second immunization; the plot on the right shows anti-PD1 IgG levels in the same set of plasma samples as in the IgG plot. The x-axis represents plasma concentration; the highest plasma concentrations of different samples were adjusted to have about 1μg/ml IgG, as defined by the IgG standard (IgG STD, black curve). The blue and red curves represent titration of from pre-or post-immune plasmas, respectively. Each curve is the result of analysis of plasma sample from one mouse. The background (green curve) was defined by buffer. **B.** ELISA detection of anti-PD1 IgG in plasma after immunization of mouse model 2. The plots are results of the same type of analysis as shown in **A**. **C.** Sorting of PD1-binding B cells from immunized mouse model 1 and cloning of 17D8 antibody or its variants. Splenic B cells from immunized mouse model 1 were first enriched for memory B cells with MACS purification kit. B220^+^IgG^+^PD1^+^ B cells, shown in the PD1^+^ gate with frequency, were sorted as single cells into 96 well plate. The HCs and LCs of the single cells were amplified with primers specific to the 17D8 V_H_/Cγ or 17D8 V_L_/Cκ. Antibodies with both 17D8 V_H_ and VL were defined as 17D8 antibodies (17D8 Ab), which were further separated into two categories, based on their CDR H3s. The pie chart shows the distribution of the three types of antibodies from the sorted single B cells; in total, 96 cells, the number at the center of the pie chart, were analyzed. **D.** Profiles of new CDR H3s in 17D8 antibody variants from mouse model 1. The logo plots show the CDR H3s, from top to bottom, in the original 17D8 antibody and related Nivolumab and three new types of novel anti-PD1 antibodies derived from 17D8. The pie chart shows the distribution of the three new types of CDR H3s. **E.** Sorting of PD1-binding B cells from immunized mouse model 2 and cloning of 17D8 antibody variants. This experiment was performed in the same manner as that shown in **C**. **F.** Profiles of new CDR H3s in 17D8 antibody variants from mouse model 2. The panel is organized in the same way as in **D**. **G.** Frequencies of amino acid changes in the 17D8 antibody heavy chains with the original CDR H3 and new CDR H3s from mouse model 1. Each dot represents one antibody heavy chain. **H.** Frequencies of amino acid changes in the heavy chains of 17D8 antibodies with new CDR H3s in mouse model 2.

For DNA sequence analyses of anti-PD1 antibodies, we used fluorophore-conjugated PD1-Fc protein as a probe to sort IgG^+^ PD1-specific splenic B cells from the immunized Mouse Model 1 (Figure 2C) and cloned paired HC and LC from individual B cells with single-cell RT-PCR (25). Because we were only interested in further studies of variants of the 17D8 antibody, we used primers specific to the 17D8 HCs and LCs for nucleotide sequence analysis of IgG^+^ PD1-specific splenic B cells isolated from both mouse models. From the spleen of one immunized PD1 Diversification Mouse Model 1, 62% of 96 sorted single B cells expressed both the 17D8 HC and LC (blue and orange portion in the pie chart, Figure 2C); the other 38% sorted B cells (grey portion in the pie chart, Figure 2C) did not yield any PCR products with the 17D8 HC-specific primer, presumably because these B cells expressed mouse IgH variable regions. Among the 62% of sorted B cells, which expressed both 17D8 HCs and LCs, 44% of the HCs had the same CDR H3 as the original 17D8 HC (orange portion in the pie chart, Figure 2C); in these cases, the joining of 17D8 V_H_ segment to the DJ_H_ segment restored the original CDR H3 of the 17D8 HC, which was then selected into the pool of IgG^+^ activated B cells by PD1 immunization. Recovery of the original 17D8 HC, subsequent to its variable region exon assembly by V(D)J recombination, confirmed that the immunization and antibody isolation procedures were highly specific for anti-PD1 antibodies. The other 18% of the HCs had CDR H3s that diverged from the 17D8 CDR H3 both in sequence and length, due to diversification at the V_H_-DJ_H_ junction (the blue portion of the pie chart, Figure 2C). Based on length and sequence, the new PD1 specific CDR H3s could be divided into three groups, which may have derived from clonal expansion of three main precursors (Figure 2D). Most of the new CDR H3s shared amino acid sequences at the N-terminal (“AR”) and C-terminal regions (“DDY”) with the original 17D8 CDR H3, because these residues were encoded by the V_H_ and the assembled DJ_H_ segments, respectively. The Asn (“N”) amino acid residue at the upstream end of the DJ_H_ segment of the 17D8 CDR H3 was prone to be altered during V_H_-DJ_H_ joining, as shown in the new CDR H3s (Figure 2D). All the LCs paired with 17D8 HCs corresponded to the pre-rearranged 17D8 LC (Figure 2C), which was expressed in most naïve B cells (Figure 1F and 1H).

We did the same DNA sequence analysis of the anti-PD1 antibodies by sorting IgG+ PD1 specific B cells from immunized Mouse Model 2 (Figure 2E). From the spleen of one immunized Mouse Model 2, 73% of 96 sorted single B cells expressed both the 17D8 HC and LC (the blue portion of the pie chart, Figure 2E). The remaining 27% of sorted B cells did not yield positive signals for 17D8 HC in single-cell PCR and presumably expressed mouse IgH variable regions (the grey portion in the pie chart, Figure 2E). In the PD1 Diversification Mouse Model 2, the 17D8 V_H_ recombined with mouse Ds and J_H_s, and all HCs had distinct CDR H3s from the 17D8 antibody. Based on CDR H3 sequence, most of the antibodies belonged to three groups (Figure 2F). Each group likely arose from clonal expansion of a founder B cell that expressed a primary antibody with the primordial CDR H3. Again, all the LCs paired with 17D8 HCs corresponded to the pre-rearranged 17D8 LC (Figure 2E).

In addition to CDR H3 diversification in PD-1 Diversification Mouse Model 1, SHM occurred throughout the reassembled 17D8 HC and the knock-in 17D8 LC variable region exons (Figure S1A and S1B, Figure 3A and Figure S2A). Overall, the PD1 Diversification Mouse Model 1 HCs with new CDR H3s had higher frequencies of amino acid changes, due to SHM, than the original 17D8 HC (Figure 2G). The difference may be attributable to the initial PD1 binding strength of their respective primary antibodies. Since the 17D8 antibody already binds strongly to PD1, SHM may not increase antigen-binding affinity substantially to confer selective advantage during the GC reaction. In contrast, primary antibodies with new CDR H3s may bind PD1 less stably than the 17D8 antibody, consequently allowing strong selection for mutations that increased PD-1 binding affinity. As a sign of antigenic selection, some of the amino acid changes (Figure 3A and Figure S2A), as well as the underlying SHMs (Figure S1A), were recurrent among different antibodies, and acquired mutations appeared more concentrated in CDRs (e.g. the N to D mutation in CDR H1 and N to K or R mutation in CDR H2, Figure 3A and Figure S1A). Like antibodies with new CDR H3s from Model 1, most of the antibodies from Model 2 contained substantial levels of SHMs (Figure S1C and S1D) and corresponding amino acid changes (Figure 2H, Figure 3B and Figure S2B). As hypothesized above, the primary antibodies for these new anti-PD1 antibodies probably interacted with PD-1 weakly, leaving ample room for affinity maturation by SHM.

**Figure 3.**
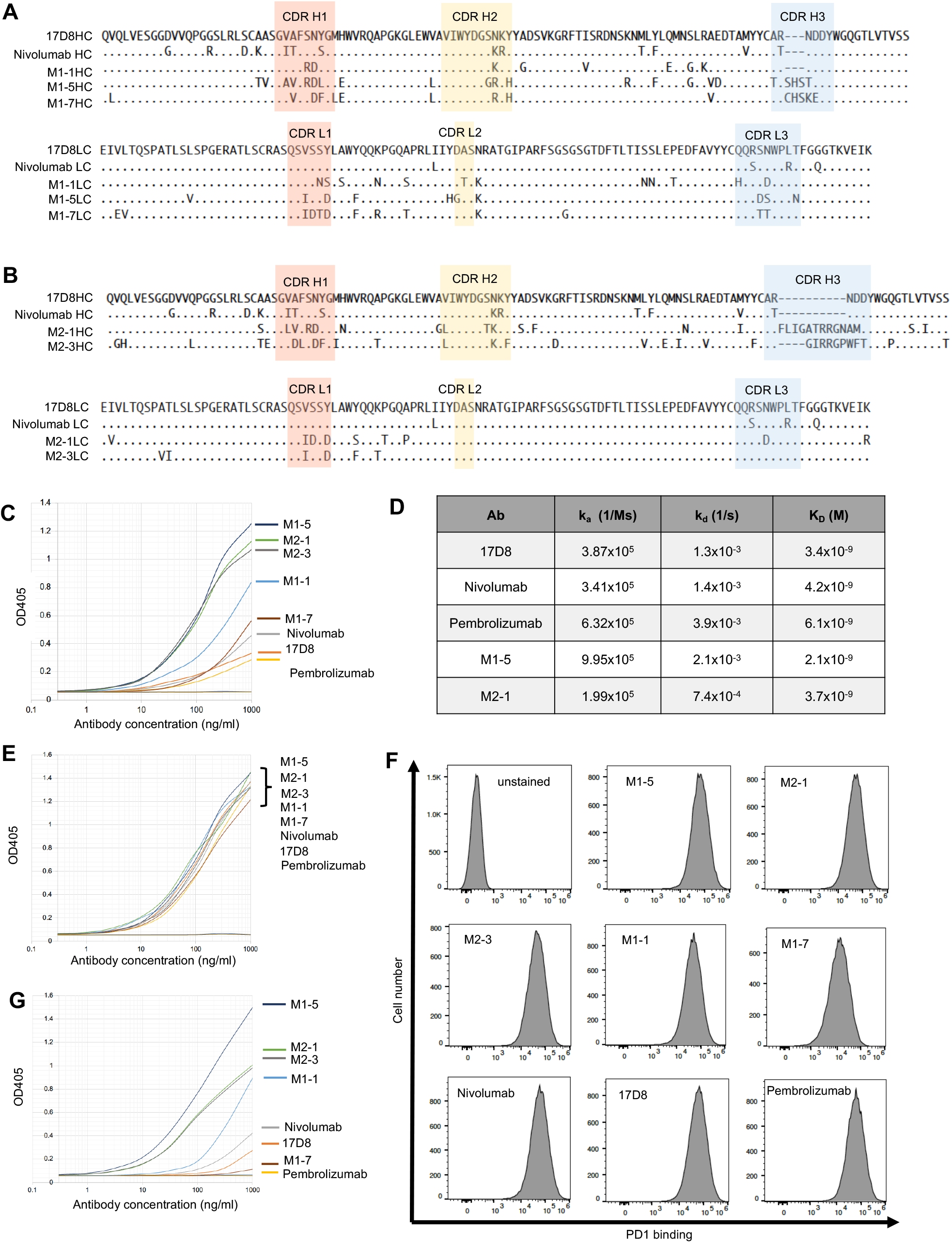
Analysis of PD1-binding activities of the 17D8 variant antibodies. **A.** Sequence comparisons of three new anti-PD1 antibodies from Mouse Model 1 with the 17D8 antibody and Nivolumab. The top and bottom parts of this section show the alignments of the amino acid sequences of IgH (HC) and IgL (LC) variable regions, respectively. In the alignment, only amino acid residues that differ from the 17D8 sequence are shown; “.” Indicates identity; “-“ represents gap in alignment. The CDRs are shaded with colors. **B.** Sequence comparisons of two new anti-PD1 antibodies from Mouse Model 2 with the 17D8 antibody and Nivolumab. The top and bottom parts of this section show the alignments of the amino acid sequences of IgH (HC) and IgL (LC) variable regions, respectively. **C.** ELISA analysis of PD1-binding activities of 17D8 antibody, Nivolumab, Pembrolizumab and five new anti-PD1 antibodies isolated from Mouse Models 1 and 2. In this ELISA experiment, PD1 extracellular domain, without Fc fusion, was coated on the ELISA plate. The x-axis represents antibody concentration in log10 scale; the y-axis displays OD405, which correlates with the levels of antibody binding to immobilized PD1. The titration curves were marked with the corresponding antibodies to the right of the plot; the order, from top to bottom, indicates the relative binding activities of the antibodies. **D.** Biacore analysis of binding kinetics of select anti-PD1 antibodies. In this assay, antibodies were immobilized on sensor chips, and PD1 extracellular domain, the same protein used in the ELISA experiment in **C**, flowed through sensor chip. The kinetics of PD1/antibody interaction was measured in real time. The table lists the association rate constant (k_a_), dissociation rate constant (k_d_) and equilibrium dissociation constant (K_D_) for each antibody. **E.** ELISA analysis of the same antibodies in Section **C**, but with PD1-Fc fusion as the coating antigen on ELISA plate. The ELISA plots are labeled the same way as in Section **C**. Since the binding curves of all the antibodies largely overlapped in this ELISA experiment, their binding activities are similar in this assay, and the order of antibodies to the right of the plot does not indicate their relative binding activities. **F.** FACS analysis of binding of anti-PD1 antibodies to PD1 expressed on cell surface. The x-axis of the FACS plots represent binding levels of the antibodies; the y-axis of the plots represents cell number. **G.**ELISA analysis of the same antibodies in Section **C**, but with PD1 N-terminal-GST fusion protein as the coating antigen on ELISA plate. The ELISA plots are labeled the same way as in Section **C**.

### Analysis of PD1 binding activities of the new anti-PD1 antibodies

To test PD1 binding activity of the new antibodies, we expressed some of the cloned HC and LC pairs as recombinant antibodies. We chose 12 antibodies representing different CDR H3 groups from the two mouse models (Figure 3A and 3B, Figure S1A-S1D, Figure S2A and S2B). We referred to the 7 antibodies isolated from PD1 Diversification Mouse Model 1 as M1-1 to M1-7 and the 5 antibodies isolated from PD1 Diversification Mouse Model 2 as M2-1 to M2-5 (Figure 3A and 3B, Figure S1A-S1D, Figure S2A and S2B). For comparison, we also produced the 17D8 antibody, Nivolumab and Pembrolizumab; the latter two antibodies are used in cancer immunotherapy (35, 36). We used ELISA to compare the PD1 binding activities of the new and previous anti-PD1 antibodies. The ELISA coating antigen was a recombinant protein of the extracellular domain of PD1, without fusion to other proteins so that the ELISA measured binding activities specific to PD1. Based on this assay, these antibodies exhibited a wide range of PD1 binding activities, and some of the new anti-PD1 antibodies (e.g. M1-5, M2-1 and M2-3 in Figure 3C; M1-4, 1-6, M2-5, M1-3, M2-2 in Figure S2C) outperformed, by far, the 17D8 antibody, Nivolumab and Pembrolizumab (Figure 3C and Figure S2C). We chose two of the strongest new anti-PD1 antibodies (M1-5 and M2-1 in Figure 3C) to quantify their PD1 binding affinity with Biacore, in side-by-side comparison with the 17D8 antibody, Nivolumab and Pembrolizumab (Figure 3D). Contrary to the ELISA results, all five antibodies showed similar PD1 binding affinities, with KD values in the nM range, which is generally considered high affinity antibody/antigen interaction. Biacore analysis is expected to be a more reliable measure of antibody binding affinity than ELISA. In the Biacore assay, PD1 was in solution phase, and all regions of the PD1 molecule should be accessible to antibodies. In contrast, immobilization of PD1 molecules on ELISA plate may occlude certain regions from antibody recognition. Under this condition, antibody binding activity will be influenced by epitope accessibility, and the apparent different PD1 binding activities of the antibodies in ELISA (Figure 3C) may, in fact, reflect distinct epitope specificities. For example, antibodies M1-5, M2-1 and M2-3 may target epitopes that are readily accessible on immobilized PD1 molecules in ELISA, whereas the PD1 epitopes for the 17D8 antibody, Nivolumab and Pembrolizumab may be partially obstructed by the ELISA plate.

To test this hypothesis, we coated ELISA plates with PD1-Fc, which was used initially to screen serum response of immunized mouse models and showed robust interaction with anti-PD1 IgG from the plasma (Figure 2A and 2B). PD1-Fc may attach to ELISA plates in a way that renders PD1 more accessible to antibody interaction. Indeed, with PD1-Fc as the coating antigen, all antibodies showed similar PD1 binding activities in ELISA (Figure 3E and Figure S2E). As another assay for PD1 binding activities, we tested the binding of these antibodies to PD1 expressed on the surface of NS1 cell, a mouse plasmacytoma cell line; this assay provides a more physiological condition for PD1 recognition than ELISA, and all regions of the PD1 extracellular domain should be readily accessible for antibody interaction. As in the PD1-Fc ELISA (Figure 3E and Figure S2E), most of the antibodies analyzed in this experiment showed comparable levels of binding to surface-expressed PD1 (Figure 3F and Figure S2F); only two antibodies, M1-7 and M1-2, showed appreciably weaker binding activity than the other antibodies (Figure 3F and Figure S2F). Thus, the results from the Biacore measurement (Figure 3D), PD1-Fc ELISA (Figure 3E and Figure S2D), and surface PD1-binding assay (Figure 3F and Figure S2F) all suggested that the new anti-PD1 antibodies have similar binding affinities to PD1 as the original 17D8 antibody. Nonetheless, these antibodies appear to recognize different PD1 epitopes, which have different levels of accessibility on immobilized PD1 molecule, and the apparent binding activities of the anti-PD1 antibodies in the PD1 ELISA experiment (Figure 3C) may actually correlate with the accessibility of their respective epitopes on immobilized PD1.

The new anti-PD1 antibodies originated from primary antibodies that had the same CDR H1 and CDR H2 in the 17D8 V_H_ segment. The structure of the 17D8 antibody and PD1 complex has not been reported. However, given the homology between the 17D8 antibody and Nivolumab, the structure of Nivolumab and PD1 complex (37) could serve as a guide to infer the roles of CDRs of 17D8 and related variant antibodies in PD1 interaction. In the case of Nivolumab, the CDR H1 and CDR H2 contact the N-terminal loop of PD1 (37). Thus, the 17D8 V_H_ containing primary antibodies in both PD1 Diversification Mouse Models may be predisposed to interact with the N-terminal loop of PD1. The main distinction between the new and the original 17D8 antibodies lie in CDR H3. The CDR H3 of Nivolumab, and potentially the homologous CDR H3 of 17D8, interacts with the FG loop of PD1 (37). The unrelated CDR H3s in the new anti-PD1 antibodies may target other regions of PD1. In some cases, the new CDR H3s may have evolved in the context of strengthening the interaction between the N-terminal loop and the CDR H1 and H2 in the 17D8 V_H_ segment; such antibodies would be focused on the N-terminal loop. Since the N-terminal loop is a linear epitope, it may be more accessible than conformational epitopes, such as the epitope for the 17D8 antibody that involves two separate regions of PD1. This difference may underlie the more robust binding activities of certain new antibodies (e.g. M1-5, M2-1 and M2-3) in PD1 ELISA than the 17D8 antibody (Figure 3C).

To test this hypothesis, we appended the N-terminal loop of PD1 to GST protein and tested the binding of the anti-PD1 antibodies to this fusion protein in ELISA. The new anti-PD1 antibodies exhibited a range of binding activities to the PD1 N-terminal loop-GST fusion (Figure 3G and Figure S1E). Some antibodies (e.g. M1-5 in Figure 3G; M1-4, M2-5, M1-6 in Figure S2E) showed strong binding activity in this ELISA, suggesting that the N-terminal loop may constitute their principle epitope. At the other end of the spectrum, some antibodies showed no detectable binding to the PD1 N-terminal loop-GST fusion protein (e.g. M1-7 in Figure 3G; M1-2 in Figure S2E), suggesting that their epitopes may involve other regions of PD1. Pembrolizumab is a representative of this type of antibody (Figure 3G), as the antibody does not contact the N-terminal loop of PD1 (38). The 17D8 antibody and Nivolumab also bound poorly to the PD1 N-terminal loop-GST fusion (Figure 3G), because these antibodies require additional regions of PD1 (e.g. the FG loop) for stable interaction (37). Between the two extremes, the other antibodies showed intermediate levels of binding to the PD1 N-terminal loop (e.g. M2-1, M2-3 and M1-1 in Figure 3G; M2-2, M1-3, M2-4 in Figure S2E); for these antibodies, the N-terminal loop may be part of their epitopes. Examining these antibodies as a whole, the binding to the PD1 N-terminal loop-GST fusion protein correlates generally with the binding to unfused PD1 in ELISA (compare Figure 3G with 3C and Figure S2E with S2C), presumably because the N-terminal loop of immobilized PD1 was readily accessible for antibody interaction. This experiment helped to resolve the discrepancies of PD1 binding activities of some antibodies in different assays (Figure 3C-3F). More importantly, the comparison of PD1 binding activities of new anti-PD1 antibodies versus the original 17D8 antibody in different assays revealed epitope diversification of the prototype in both Mouse Models.

### Stimulation of PD1/PD-L1 interaction by new anti-PD1 antibodies

Antibodies targeting different PD1 epitopes may exert distinct effects on PD1 interaction with its ligands. We used a cell-based assay to test this possibility. In this experiment, we measured the binding of the two PD1 ligands, PD-L1 (13, 39) or PD-L2 (14, 40), to PD-1 expressed on the NS1 cell surface. Although both PD-L1 and PD-L2 bound to surface expressed PD1 in this assay, PD-L1 exhibited lower binding activity than PD-L2 (compare PD-L1 in Figure 4A with PD-L2 in Figure 4B), in agreement with biochemical measurements of PD1/ligand interaction (41). As expected from its homology to Nivolumab, addition of the 17D8 antibody abrogated PD1 interaction with PD-L1 (Figure 4A) or PD-L2 (Figure 4B). One new anti-PD1 antibody, M1-1, had the same inhibitory effect on PD1/ligand interaction (Figure 4A and 4B). By contrast, the other 4 new antibodies (M2-1, M1-5, M1-7 and M2-3) enhanced PD1/PD-L1 interaction to varying degrees, with antibodies M1-7 and M2-3 exhibiting the strongest effects (Figure 4A). On the other hand, these 4 antibodies (M2-1, M1-5, M1-7 and M2-3) did not affect PD1/PD-L2 interaction (Figure 4B). Additional new anti-PD1 antibodies exhibited similar activities: enhancing PD1/PD-L1 interaction but having no effect on PD1/PD-L2 interaction (Figure S3A and S3B). One new antibody, M2-4, inhibited PD1/PD-L1 interaction, but had minimal effect on PD1/PD-L2 interaction (Figure S3A and S3B). These results showed that diversification of the 17D8 antibody produced variants with different activities: antibody M1-1 can inhibit both PD1/PD-L1 (Figure 4A) and PD1/PD-L2 (Figure 4B) interaction, antibody M2-4 can block PD1/PD-L1 (Figure S3A) but not PD1/PD-L2 (Figure S3B) interaction, and the other 10 antibodies can stimulate PD1/PD-L1 interaction to varying degrees (Figure 4A and S3A) but do not affect PD1/PD-L2 (Figure 4B and S3B) interaction.

**Figure 4.**
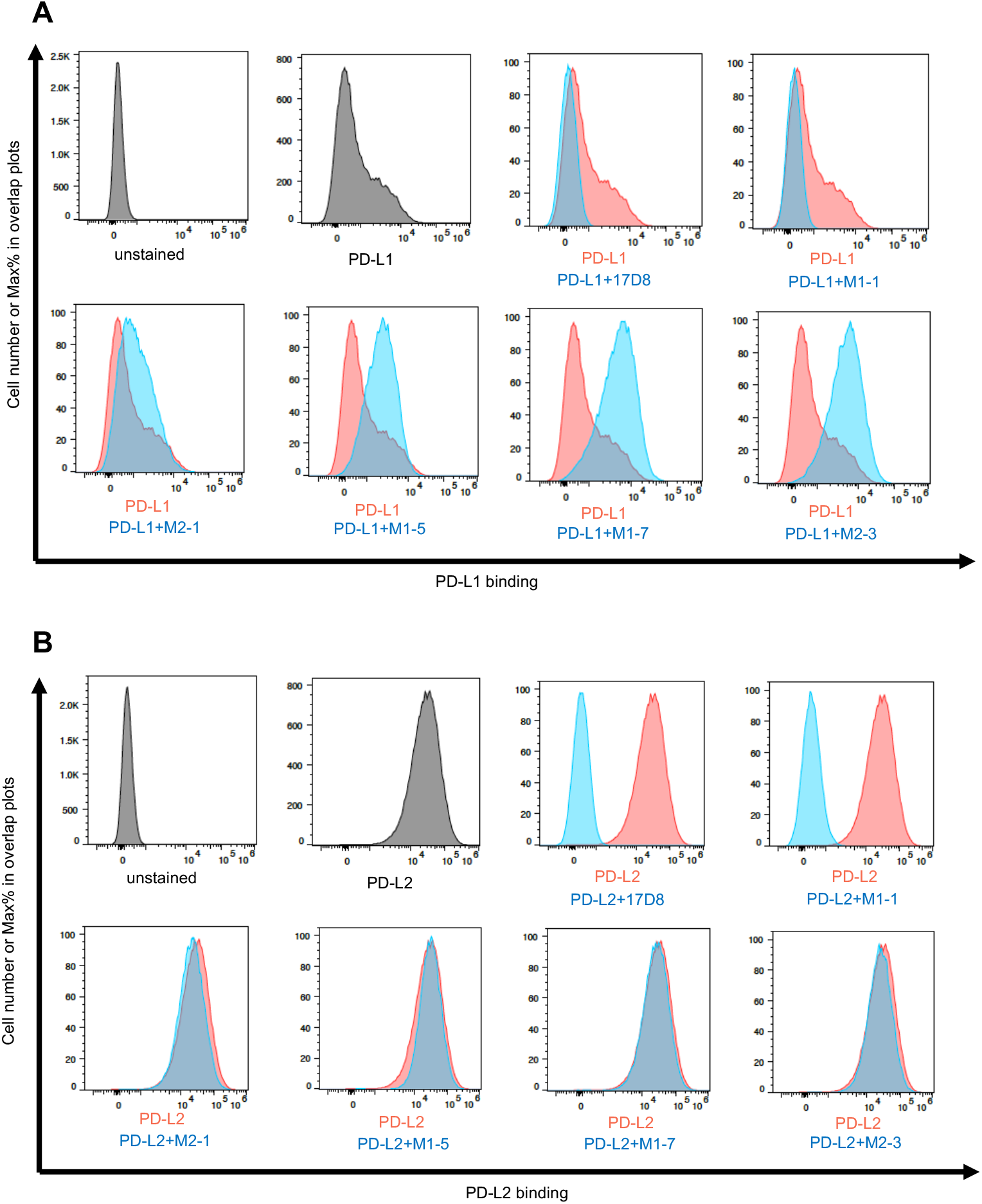
The effects of anti-PD1 antibodies on PD1/ligand interaction. **A.** FACS analysis of the effects of anti-PD1 antibodies on PD1/PD-L1 interaction. The x-axis of the plots represents the levels of PD-L1 binding to PD1 expressed on NS1 cell surface; the y-axis represents relative cell number, with the highest peak set at 100% (modal mode). The addition of PD-L1 and various anti-PD1 antibodies are indicated underneath the plots. In the plot “PD-L1”, only PD-L1 was added to the binding reaction, and this data was used in all the subsequent overlay plots, as represented by the red histogram; the blue histograms show PD-L1 binding to PD1 in the presence of various antibodies. **B.** FACS analysis of the effects of anti-PD1 antibodies on PD1/PD-L2 interaction. The plots in this section is labeled in the same manner as in **A**.

PD1 blocking antibodies work by a competition mechanism. Based on structural studies, Nivolumab binding to PD1 clashes with PD1/ligand interaction, and the homologous 17D8 antibody likely functions similarly. As shown above, the new anti-PD1 antibody M1-1 also blocked PD1/ligand interaction. Among the 12 new antibodies analyzed in the PD1/ligand interaction assay (Figure 4 and Figure S3), the M1-1 antibody was the only one that preserved the CDR H3 of the 17D8 antibody (Figure 3A). Because of the CDR H3 conservation, the M1-1 antibody may interact with PD1 in a similar manner as the 17D8 antibody to inhibit PD1/ligand interaction. The other new anti-PD1 antibodies contained different CDR H3s from the 17D8 antibody (Figure 3A and 3B, Figure S1A-S1D, Figure S2A and S2B). In addition, some of these new antibodies also accumulated substantial levels of SHM throughout both the IgH and IgL variable regions (Figure 3A and 3B, Figure S1A-S1D, Figure S2A and S2B). Altogether, these sequence changes, especially those in CDRs, could alter the way these antibodies contact PD1; for example, some of the new antibodies (e.g. M1-5, M2-1 and M2-3 in Figure 3G) targeted primarily the N-terminal loop of PD1, whereas the 17D8 antibody contacted additional regions of PD1. The N-terminal loop specificity cannot account for the stimulatory effect of the new anti-PD1 antibodies. One of the best stimulatory antibodies, M1-7 (Figure 4A), barely associated with the PD1 N-terminal loop in ELISA (Figure 3G); by contrast, the M2-1 antibody bound strongly to the PD1 N-terminal loop (Figure 3G), but this antibody had minimal stimulatory effect on PD1/PD-L1 interaction (Figure 4A). Structural studies of the antibody/PD1/PD-L1 complex would help to elucidate the mechanism that underlies the stimulatory effect of some of the new anti-PD1 antibodies. In such structural studies, comparison of the M1-7 and M2-3 antibodies could be particularly informative, as these two antibodies showed comparable stimulatory effects on PD1/PD-L1 interaction; but they appeared to target different epitopes in PD1, based on their distinct binding activities to PD1 and PD1 N-terminal loop in ELISA. Another unresolved question is why, with the exception of the M1-1 antibody, none of the new antibodies inhibit PD1/PD-L2 interaction. PD-L1 and PD-L2 bind to PD1 in a similar fashion (41); however, PD-L2 has higher affinity for PD1 than PD-L1 (41) perhaps rendering the PD1/PD-L2 association less susceptible to modulation by antibodies.

In summary, this study served as a proof-of-concept validation of our antibody diversification strategy to derive variant antibodies from a prototype. Moreover, this study yielded a panel of new anti-PD1 antibodies that deserve further investigation for potential therapeutic applications. For example, among the new anti-PD1 antibodies, M1-7 and M2-3 exerted the most obvious stimulatory effects on PD1/PD-L1 interaction (Figure 4A). If these *in vitro* stimulatory effects are translatable to *in vivo* situations, these antibodies (M1-7 and M2-3) could potentiate PD1-dependent inhibitory pathway to suppress deleterious T cell activities in autoimmune diseases. In this regard, PD1/PD-L1 interaction plays an important role in preventing autoimmunity (42–52). These antibodies do not affect PD1/PD-L2 interaction. PD-L2 is primarily expressed on professional antigen presenting cells, whereas PD-L1 has a broader expression pattern (53). Thus, these antibodies may not affect T cell priming by professional antigen-presenting cells, and their primary impacts would be on effector T cell activity toward PD-L1^+^ target cells. The specificity could be exploited to dampen autoimmune T cell attack on host tissues, without general immune suppression. These antibodies could also inhibit T cell activity independently of PD-L1. IgG isoforms of the antibodies can attach to cell surface via Fcγ receptors (54) and engage PD1 directly on T cells to down-regulate T cell activity. All these potential applications will require functional evaluation *in vivo*.

## Materials and Methods

### Generation and characterization of mouse models to diversify the 17D8 antibody

The genetic modifications for the mouse models were introduced into mouse embryonic stem (ES) cells. For replacement of the mouse V_H_81X segment with the 17D8 V_H_ segment, we generated a homologous recombination construct, which contains homology arms that flank the mouse V_H_81X segment. We transfected the homologous recombination construct into a previously established ES cell line that was used to generate a mouse model for testing HIV-1 vaccine candidates. In this ES cell line, the mouse V_H_81X segment was replaced with the human V_H_1-2 gene segment. Moreover, the IGCRI regulatory element was deleted to increase the usage of the human V_H_1-2 or any human gene segment in place of the mouse V_H_81X segment. After transfection, stable genomic integration of the recombination construct into ES cells was selected with G418, based on a neomycin resistance gene in the construct. The stable clones were screened, with Southern hybridization, for correct integration of the human 17D8 V_H_ segment into the V_H_81X locus. The neomycin resistance gene was removed via deletion by flanking loxP sites by transduction of the ES clone with Adenovirus-cre. All the genetic modifications involved in the mouse models, including 17D8 DJ_H_ and 17D8 LC KIs, were introduced into the respective loci, as diagramed in Figure 1A and 1B, in the same manner. The ES clones with the complete set of genetic modifications for mouse model 1 or model 2 were injected into Rag2 deficient blastocysts to generate chimeric mice. All the immunization experiments in this study were carried out with such chimeric mice.

To determine B cell reconstitution in chimeric mice, blood samples were collected from the chimeric mice. B cells were stained with PE-anti-B220 antibody and FITC-anti-IgM antibody (Figure 1C and 1D). The FACS reaction was analyzed on a BD FACS Calibur machine, and the data were plotted with FlowJo software. To analyze the representation of 17D8 HC among B cells in the chimeric mice (Figure 1E and 1G), we fused splenic B cells from the mouse models with NS1 plasmacytoma cells to generate hybridomas. V(D)J recombination products involving the 17D8 V_H_ segment was detected by PCR amplification with a primer specific to the 17D8 V_H_ segment and a primer downstream of the J_H_ region. To verify the expression of 17D8 LC in B cells, we purified RNA from splenocytes of the mouse models. 5’RACE for total Igκ cDNA was initiated with a primer specific to Cκ, and the PCR products were cloned into shuttle vector. Individual clones were sequenced, and all corresponded to the 17D8 LC (Figure 1F and 1H).

### Immunization of mouse models and isolation of anti-PD1 antibody genes

The immunogen, PD1-GST fusion protein, consisted of the extracellular domain of human PD1 fused at the C-terminus to GST. The fusion protein was expressed in E. coli and purified with glutathione agarose. Each mouse was immunized with 50μg of PD1-GST protein plus Inject-Alum by intraperitoneal injection. The mouse model 1, from which the 319-9-x antibodies were isolated, was immunized 5 times altogether; the mouse model 2, from which the 397-27-x antibodies were isolated, was immunized 4 times in total.

To monitor the induction of anti-PD1 antibody by immunization, blood samples were collected before and after the second immunization. Plasma was separated from the cellular fraction by centrifugation. IgG concentrations in the plasma was measured with ELISA; in this assay, unlabeled goat anti-mouse IgG antibody was used for capture, alkaline phosphatase (AP) conjugated goat anti-mouse IgG served as secondary antibody for detection, and purified mouse IgG was the standard for quantification. Based on this initial measurement, the plasma concentrations were adjusted to the same IgG concentrations, which were verified in ELISA (Figure 2A and 2B). The levels of anti-PD1 IgG in the same plasma samples were measured with ELISA (Figure 2A and 2B). For ELISA detection of anti-PD1 antibodies (Figure 2A and 2B), ELISA plates were coated with PD1-Fc fusion protein, which is composed of the extracellular domain of human PD1 and the human IgG4 Fc region. The use of PD1-Fc fusion avoided detection of anti-GST antibodies that were induced by PD1-GST immunization. The PD1-Fc fusion was expressed in 293F cells and purified via a C-terminal His tag on Ni-column. The secondary antibody for the ELISA was AP-goat anti-mouse IgG (Figure 2A and 2B).

To isolate PD1-specific B cells, spleen was dissected 5 days after the last immunization. IgG^+^ B cells were enriched with a memory B cell purification kit. The IgG^+^ B cell preparation was stained with Alexa488-PD1-mFc, PE-anti-B220 and Sytox blue, and the B cells were already stained with APC-anti-IgG antibodies from the memory B cell purification kit. PD1-mFc fusion protein consisted of the extracellular domain of human PD1 and mouse IgG1 Fc region. The fusion protein was expressed in 293F cells, purified via C-terminal His tag on Ni-column and labeled with Alexa488. The use of PD1-mFc fusion selected for PD1-specific B cells, instead of GST-specific B cells, which should also be present in the IgG^+^ B cell population. PD1-specific IgG+ B cells were sorted as single cells into 96 well plates on FACS Arias (Figure 2C and 2D). The Ig HCs and LCs of the single cells were amplified with primers specific for the 17D8 V_H_/mouse Cγ1, 2a, 2b or 17D8 Vκ/mouse Cκ, respectively. The cells that were positive for both the 17D8 HC and LC PCRs were counted as expressing 17D8 related antibodies, as shown in the pie chart (Figure 2C and 2D). The PCR products for the HC or the LC were sequenced, and their sequences were aligned with the 17D8 HC or LC sequences (Figure S1) with MegaAllign software, and mutation frequency (Figure 2G and 2H) was based on substitution ratios of the V_H_ segment. CDR H3 logo plots (Figure 2E ad 2F) were generated with WebLogo.

### Analysis of the binding activities of anti-PD1 antibodies

For the ELISA experiment in Figure 3A, anti-PD1 antibodies were produced by transient transfection of expression constructs into 293F cells or Expi293 cells; all antibodies were expressed as human IgG4/κ antibodies. These antibodies contained a 6xHis tag at the C-terminus of the Fc region and were purified from culture supernatant on Ni-column. For measurement of PD1 binding activity, ELISA plates were coated with PD1-Fc fusion protein. Antibodies were added to the plates; in the dilution series, the highest concentration of all the antibodies were 1μg/ml, as confirmed by ELISA quantification of IgG concentration. After washing, antibodies retained on the plates were detected with AP-anti-human kappa antibody. For the ELISA experiment in Figure 5, ELISA plates were coated with PD1 N-terminal loop-GST fusion protein; the N-terminal loop peptide is: LDSPDRPWNP. The N-terminal loop-GST protein was produced in E. coli and purified on glutathione agarose. The same anti-PD1 antibodies as those used for the PD1-Fc fusion ELISA was analyzed in the PD1 N-terminal loop-GST ELISA, and AP-anti-human kappa antibody served as the secondary antibody for detection.

To test the binding of anti-PD1 antibodies to surface expressed PD1, we expressed full-length human PD1 in mouse plasmacytoma cell line, NS1. As confirmed by FACS staining, the parental NS1 cells did not exhibit appreciable cross-reactivity with any of the anti-PD1 antibodies in this study (Figure S2). We transfected into NS1 cells a construct that expresses the full length human PD1 and obtained clones with stable integration of the expression construct. We chose one clone, referred to as PD1-NS1, for the FACS analysis in Figure 3B and Figure 4. For the FACS analysis in Figure 3B, the PD1-NS1 cells were incubated with anti-PD1 antibodies. The binding of the PD1 antibodies were revealed with PE-anti-human kappa antibody. The FACS staining reaction was analyzed on Attune NXT FACS machine, and the data were plotted with FlowJo 10 software.

Biacore analysis of the kinetics of the interaction between anti-PD1 antibodies and PD1 was performed by Affina Biotechnologies. Given the quantitative and sensitive nature of the assay, we tried to minimize non-physiological modifications of the antibodies and PD1. Thus, the antibodies did not contain C-terminal His tag; these antibodies were expressed as human IgG4/k antibodies in Expi293 cells and purified on Protein A column. The PD1-Fc fusion protein contains a TEV protease cleavage site. After TEV digest, the PD1 portion was separated from the Fc region. Under physiological conditions, PD1 exists as a monomer. However, in the context of the PD-Fc fusion, PD1 is dimerized through the Fc region and may engage in bivalent interactions with IgG antibodies. The avidity effect could cause major deviations from the actual binding affinity between monomeric PD1 and single antigen-binding site. Separation of PD1 from the Fc portion prevented this error. In the biacore assay, the antibodies were immobilized on sensor chips and PD1 was passed over the immobilized antibodies. Association and dissociation of antibodies and PD1 were detected in real time. Based on these data, the on-rate and off-rate of antibody/PD1 interaction were determined, and the KD value was derived by dividing the off-rate with on-rate.

### Analysis of the effects of anti-PD1 antibodies on PD1/ligand interaction

PD-L1 and PD-L2 were expressed as fusion proteins with human IgG4 Fc regions in Expi293 cells, purified from the culture supernatant via C-terminal His tag on Ni-column and biotinylated on Avi-tag at the C-terminus. For PD1/ligand interaction experiments in Figure 4, the biotinylated PD-L1 or PD-L2 were incubated with PD1-NS1 cells with or without anti-PD1 antibodies. After washing, PD-L1 or PD-L2 retained on cell surface were revealed with PE-Streptavidin. The binding reaction was analyzed on Attune NXT flow cytometer, and the data were plotted with Flowjo software.

## Acknowledgements

We thank Peiyi Huang for microinjection of the ES cells to generate RDBC chimeric mice. This work was supported by a Technology Development Fund from the Boston Children’s Hospital and the Howard Hughes Medical Institute (HHMI). FA is an investigator of the HHMI.

## Supplementary Materials

### Supplementary Methods

#### The Ig variable sequences of the new anti-PD1 antibodies that were used in the study

The sequences encode the Ig variable regions of the new anti-PD1 antibodies that were used in the experiments in Figure 3–4, Figure S2 and S3. In the expression constructs, the human V_H_3-33 leader sequence and the human Vκ3-11 leader sequence were appended to the 5’ end of the variable region sequences of the HCs and LCs, respectively; human V_H_3-33 and Vκ3-11 segments are germline components of the 17D8 antibody heavy and light chains, respectively. The IgH variable regions were expressed in association with the human IgG4 constant region, and the IgL variable regions were expressed in association with the human kappa constant region. The complete cDNAs for the HCs and LCs were cloned into the pcDNA expression vector and expressed in 293F or Expi293 cells.

##### M1-1HC

CAGGTGCAGTTGGTGGAGTCTGGGGGAGACGTGGTCCAGCCTGGGGGGTCCCTGAGACTCTCCTGTGCAGCGTCTGGAGTCGCCTTCAGGGACTATGGCATGCACTGGGTCCGCCAGGCACCAGGCAAGGGGCTGGAGTGGGTGGCAGTTATATGGTATGATGGAAGTAAGAAATATTATGGAGACTCCGTGAAGGGCCGATTCACCGTCTCCAGAGACAATTCCAAGAACATGTTGTATCTGGAAATGAACGGCCTGAAAGCCGAGGACACGGCAATGTATTATTGTGCGAGGAACGATGACTACTGGGGCCAGGGAACCCTGGTCACCGTCTCCTCAG

##### M1-2HC

CAGGTGCAGCTGGTGGAGTCTGGGGGAGACGTGGTCCAGCCGGGGGGGTCCCTGAGACTCTCCTGTGCGGCGTCTGGAGTCGCCTTCAGTGACTATGGCATGGAATGGGTCCGCCAGGCTCCAGGCAAGGGGCTGGAGTGGCTGGCAGTTATCTGGTATGATGGAAGTAGGAAACACTATGCAGACTCCGTGAAGGGCCGATTCACCATCTCCCGCGACAATTCCAAGAACATTCTGTATCTACAAATGAACAGCCTGAGAGCCGAGGACACGGCTATGTATTACTGTGCGAGATGCCACTCTAAAGATGACTACTGGGGCCAGGGAACCCTGGTCACCGTCTCCTCAG

##### M1-3HC

CAGGAGTGGTTGGTGGAGTCGGGGGGAGACGTGGTCCAGCCGGGGGGGTCCCTGAGACTCTCCTGTGCGACGTCTAAAGTCACCTTCAATGACTTTGGCATTCACTGGGTCCGCCAGGCTCCAGGCAAGGGACTGGAGTGGGTGGCAATTATTTGGTATGATGGAAGCAGGAATCACTACGCAGACTCCGTGAGGGGCCGATTCACCCTCTCCAGGGACAATTCCAAAAACATGGTCCATCTTCACATGAGTAGCCTGAGAACCGAGGACACGGCTATGTATTATTGTGCGCGAGGAATACACTCTAACGATGACTATTGGGGCCAGGGAACCATGGTCACCGTCTCCTCAG

##### M1-4HC

CAGGTTCAGCTGGTGGAATCTGGGGGAGACGTGGTCCAGCCGGGGGGGTCCCTGAGACTCTCCTGTGCAGTGTCTGGAGTCACCTTCGGTGACTTTGGCTTCGAATGGGTCCGCCAGGCTCCAGGCAAGGGTCTGGAGTGGGTGGCAGTTATTTGGTACGACGGAAGCAGAAAACATTATGCAGAGTCCGTGAGGGGCCGATTTACCATCTCCAGAGACAATTCCAGGAACATGATGTATCTGGAAATGACTGGACTGAGAGTCGAGGACACGGCTAAATATTACTGTACGAGAAGTCACTCTCACGAGGACTACTGGGGCCAGGGAACCCTGGTCACCGTCTCCTCAG

##### M1-5HC

CAGGTGCAGCTGGTGGAGTCTGGGGGAGACGTGGTCCAGCCTGGGGGGTCCCTGAGACTCTCCTGTACAGTGTCTGGAGCCGTCTTCCGTGACTTGGGCATGGAATGGGTCCGTCAGGCCCCAGGCAAGGGGCTGGAATGGTTGGCAGTTATATGGTACGACGGAGGTAGAAAACACTATGCGGACTCCGTGAAGGGCCGGTTCACCATCTCCAGAGACAATTCCAGGAACATGCTCTTTCTGCAAATGAATGGACTGAGAGTCGACGACACGGCTATGTATTACTGTACGAGAAGCCACTCTACCGATGATTACTGGGGCCAGGGAACCCTGGTCACCGTCTCCTCAG

##### M1-6HC

CAGGTGCAGTTGGTGGAGTCTGGGGGAGACGTGGTCCAGTCGGGGGGGTCCCTGAGACTCTCCTGTGCAGTGTCTGGAGTCGTCTTCAGTGATTATGGCTTCGAATGGGTCCGCCAGGCTCCAGGCAAGGGGCTGGAGTGGGTGGCAGTTATATGGTACGACGGAAGTAGGAAACATTATGCAGACTCCGTCCAGGGCCGATTCACCATTTCCAGAGACAATTTCCGGAACATGTTGTATCTACAAATGACTGGACTGAGAGTCGAGGACACGGCTAAATACTATTGTACGAGAAGCCACTCTCACGAGGACTACTGGGGCCAGGGAACCCTGGTCACCGTCTCCTCAG

##### M1-7HC

CAATTACAACTGGTGGAGTCTGGGGGAGACGTGGTCCAGCCTGGGGGGTCCCTGAGACTCTCCTGTGCGGCGTCTGGGGTCGTCTTCAGTGACTTTGGCCTGGAATGGGTCCGCCAGGCTCCAGGCAAGGGGCTGGAGTGGCTGGCAGTTATCTGGTATGATGGAAGTCGGAAACATTATGCAGACTCCGTGAAGGGCCGATTCACCATCTCCAGGGACAATTCCAAGAACATGCTCTATCTGCAAATGAACAGTCTGAGAGTCGAGGACACGGCTATGTACTACTGTGCGCGATGTCACTCTAAAGAGGACTACTGGGGCCAGGGAACCCTTGTCACCGTCTCCTCAG

##### M2-1HC

CAGGTGCAGCTGGTGGAGTCTGGGGGAGACGTGGTCCAGCCTGGGGGGTCCCTGAGACTCTCCTGTTCAGCGTCTGGACTCGTATTCAGAGACTATGGCATGAACTGGGTCCGCCAGGCTCCAGGCAAGGGGCTGGAGTGGGTCGGACTTATATGGTATGATGGAACTAAAAAATATTATTCAGACTTCGTGAAGGGCCGATTCACCATCTCCAGAGACAATTCCAAGAACATGTTGTATCTACAAATGAACAACCTGAGAGCCGAGGACACGGCTATTTATTACTGTGCGAGATTTCTAATAGGTGCGACGAGGAGGGGCAATGCTATGGACTACTGGGGTCAAGGGACCTCAGTCATCGTCTCATCAG

##### M2-2HC

CAGGTGCAGTTGGTGGAGTCTGGGGGAGACGTGGTCCAGCCTGGGGGGTCCCTGAGACTCTCCTGTTCAGCGTCTGGGCTCGTAATCAGTGACTATGGCATGAACTGGGTCCGCCAGGCTCCAGGCAAGGGGCTGGAGTGGGTCGGACTTATATGGTATGATGGAAGTAAAAAATATTACTCAGACTTCGTGAAGGGCCGATTCACCATCTCCAGAGACAATTCCAAGAACATATTGTATCTACAAATGAACAACCTGAGAGCCGAGGACACGGCTATGTATTACTGTGCGAGATTTCTAATAGGTGCGACGAGGAGGGGCAATGCTATGGACTATTGGGGTCAAGGAACCTCAGTCACCGTCTCATCAG

##### M2-3HC

CAGGGGCACCTGGTGGAGTCTGGGGGAGACGTGGTCCTGCCTGGGGGGTCCCTGAGACTCTCCTGTACAGAGTCTGGTGTCGACCTCAGTGACTTTGGCATACATTGGGTCCGCCAGACTCCAGGCAAGGGTCTGGAGTGGGTGGCACTTATCTGGTATGATGGAAGTAAAAAATTTTATGCTGACTCCGTGAAGGACCGATTCACCATTTCCAGAGACAATTCCAAGAATATGGTGTATCTGGAAATGATCAGCCTGAGAGTCGAGGATACGGCTATGTACTTCTGTGCGAGAGGGATACGACGGGGGCCCTGGTTCACTTACTGGGGCCCAGGGACTCTGGTTACAGTCTCTACAG

##### M2-4HC

CAGGTGCAGTTGGTGGAGTCTGGGGGAGACGTGGTCCAGCCGGGGGGGTCCCTGAGACTCTCCTGTGCAGCGTCTGGAGTCGCCTTCAGGAACTATGGCATGCACTGGGTCCGCCAGGCTCCAGGCAAGGGCCTGGAGTGGGTGGCAATTATATGGTATGATGGAAGTAATAAATATTATGCAGACTCCGTGAAGGGCCGCTTCACCATCTCCAGAGACAATTCCAAGAATATGTTGTATCTTCAAATGAATAGCCTGAGAGCCGAGGACACGGCTATGTATTACTGTGCGAGACTCTCTATAGGTACGACCCATTACTTTGATACGGACGACTACTGGGGTCAAGGAACCTCAGTCACCGTCTCCTCAG

##### M2-5HC

CAGGTGCAACTGGTGGAGTCTGGGGGAGACGTGGTCCAGCCGGGGGGGTCCCTGAGACTCTCCTGTGCAGCGTCTGGAGTCGTCTTCAGTGACTATGGCTTGTATTGGGTCCGCCAGGCTCCAGGCAAGGGCCTGGAGTGGGTGGCCCTTATATGGTATGATGGGAGTAAGAAATTTTATGCTGACTCCGTGAAGGGCCGATTCTCCATCTCCAGAGACAATTCCAAGAACATGTTGTATTTACAAATGAATAATTTGAGAGCCGACGACTCGGCTATTTATTATTGTTCGAGAGGGATACGACAGGGGCCATGGTTTGCTTACTGGGGCCAAGGGACTCGTGTCACTGTCTCTCCAG

##### M1-1LC

GAAATTGTGTTGACACAGTCTCCAGCCACCCTGTCTTTGTCTCCAGGGGAAAGAGCCACCCTCTCCTGCAGGGCCAGTCAGAGTGTTAGCAACTCCTTATCCTGGTACCAACAGAACCCTGGCCAGTCTCCCAGGCTCATCATCTATGATACATCCAAGAGGGCCACTGGCATCCCAGCCAGGTTCAGTGGCAGTGGATCTGGGACAGACTTCACTCTCACCATCAACAATCTAGAGACTGAAGATTTTGCAGTTTATTACTGTCACCAGCGTAGCGACTGGCCTCTCACTTTCGGCGGAGGGACCAAGGTGGAGATCAAAC

##### M1-2LC

GAAATTGTGTTGACACAGTTTCCGGCCACCCTGTCTCTGTCTCCCGGGGAAAGAGCCACCCTCTCCTGCAGGACCAGTCAGAATATTGACAGCGACTTAGCCTGGTTCCAACAGAAACCTGGCCAGGCTCCCAGGCTCATCATCTATGATGCATCCAACAGGGCCACTGGCATCCCAGCCAGGTTCAGTGGCGGTGGGTCTGGGACAGACTTCACTCTCACCATCACCAGCCTAGAGCCTGAAGATTTTGCAGTTTATTACTGTCAGCAGCGTACCACCTGGCCTCTCACTTTCGGCGGAGGGACCAAGGTGGAGATCAAAC

##### M1-3LC

GAAATTGTGTTGACACAGTCTCCAGCCACCCTGTCTTTGTCTCCAGGGGAAAGAGCCACCCTCTCCTGCCGGACCAGTCAGAGTGTTAGCAGCGACTTAGCCTGGTTCCAACAGAAACCTGGCCAGGCTCCCAGGCTCTTCATATTTGATGCATCCAAAAGGGTCAATGGCATCCCAGCCAGGTTCAGTGGCAGTGGGTCTGGGACAGACTTCACTCTCACCATCAGCAGCCTGGAACCTGAAGATTTTGCAGTTTATTATTGTCAGCAACGTACCGACTGGCCTCTCACTTTCGGCGGAGGGTCCAGGGTGGAGATCAAAC

##### M1-4LC

GAAATTGTGTTGACACAGTCTCCAGCCACCCTGTCTTTGTCTCCAGGGGAAAGAGCCACCCTCTCCTGTAGGGCCAGTCAGAGTATTAGCAACTACTTAGCCTGGTTCCAACAGAAATCTGGCCAGGCTCCCAGGCTCATCATCCATGATGCATTTAAACGGGCCGCTGGCATCCCAACCAGGTTCAGTGGCAGTGGGTCTGGGACAGACTTCACTCTCACCATCAGCAGTCTAGAGCCTGAAGATTTTGCAGTTTATTATTGTCAGCAGCGTGACAACTGGCCTCTCAATTTCGGCGGAGGGACTAAGGTGGAGATCAAAC

##### M1-5LC

GAAATTGTGTTGACACAGTCGCCAGCCACTCTGTCTGTGTCTCCAGGGGAAAGAGCCACCCTCTCCTGTAGGGCCAGTCAGAGTATTAGCAGCGACTTAGCCTGGTTCCAACAGAAACCTGGCCAGGCTCCCAGGCTCATCATCCATGGTGCATCCAAAAGGGCCACTGGCATCCCAGCCAGGTTCAGTGGCAGTGGGTCTGGGACAGACTTCACTCTCACCATCAGCAGCCTAGAGCCTGAAGATTTTGCGGTTTATTACTGTCAGCAGCGTGACAGCTGGCCTCTCAATTTCGGCGGAGGGACCAAGGTGGAGATCAAAC

##### M1-6LC

GAAATTGTGTTGACACAGTCTCCAGCCACCCTGTCTTTGTCTCCAGGGGAAAGAGCCACCCTCTCCTGTGGGGCCAGTCAGAATATTGACAACTCCTTAGCCTGGTTCCAACAGAAACCTGGCCAGGCTCCCAGGCTCATCATCTATGATGCATCTAAAAGGGCCACTGGCATCCCAGCCAGGTTCAGTGGCAGTGGGTCTGGGACAGACTTCACTCTCACCATCAGCACTCTAGAGCCTGAAGATTTTGCAGTTTATTATTGTCAGCAGCGTGACCATTGGCCTCTCAATTTCGGCGGAGGGACCAAGGTGGAAGTCAAAC

##### M1-7LC

GAAATTGAAGTGACACAGTCTCCGGCCACCCTGTCCTTGTCTCCAGGGGAAAGAGCCACCCTCTCCTGTAGGGCCAGTCAGAGTATTGACACCGACTTAGCCTGGTTCCAGCAGAGACCTGGCCAGACTCCCAGACTCATCATCTATGATGCATCCAAAAGGGCCACTGGCATCCCAGCCAGGTTCAGTGGCGGTGGGTCTGGGACAGACTTCACTCTCACCATCAGCAGCCTAGAGCCTGAAGATTTTGCAGTTTACTACTGTCAGCAGCGTACCACCTGGCCTCTCACTTTCGGCGGAGGGACCAAGGTGGAGATCAAAC

##### M2-1LC

GAAGTTGTGTTGACACAGTCTCCAGCCACCCTGTCTTTGTCTCCAGGGGAAAGAGCCACCCTCTCCTGCAGGGCCAGTCAGAGTATTGACAGCGACTTAGCCTGGTCCCAACAGAAAACTGGCCAGCCTCCCAGACTCATCATCTATGATGCATCCAACAGGGCCACTGGCATCCCAGCCAGGTTCAGTGGCAGTGGGTCTGGGACAGACTTCACTCTCACCATCAGTAGTCTAGAGCCTGAAGATTTTGCAGTTTATTATTGTCAGCAACGTAGCGACTGGCCTCTCACTTTCGGCGGAGGGACCAAGGTGGAGATCAGAC

##### M2-2LC

GAAGTTGTGTTGACACAGTCTCCAGCCACCCTGTCTTTGTCTCCAGGGGAAAGAGCCACCCTCTCCTGCAGGGCCAGTCAGAGTATTGACAGCGACTTAGCCTGGTCCCAACAGAAACCTGGCCAGCCTCCCAGACTCATCATCTATGATGCATCCAACAGGGCCACTGGCATCCCAGCCAGGTTCAGTGGCAGTGGGTCTGGGACAGACTTCACTCTCACCATCAGCAGCCTAGAGCCTGAAGATTTTGCAGTTTATTATTGTCAGCAACGTAGCGACTGGCCTCTCACTTTCGGCGGAGGGACCAAGGTGGAGATCAGAC

##### M2-3LC

GAAATTGTGTTGACACAGTCTCCAGTCATCCTGTCTTTGTCTCCAGGGGAAAGAGCCACCCTCTCCTGCAGGGCCAGTCAGAGTATTAGCAGCGACTTGGCCTGGTTCCAACAGACACCTGGCCAGGCTCCCAGGCTCATCATCTATGATGCATCCAACAGGGCCACTGGCATCCCAGCCAGGTTCAGTGGCAGTGGGTCTGGGACAGACTTCACTCTCACCATCAGCAGCCTAGAGCCTGAAGATTTTGCAGTTTATTACTGTCAGCAGCGGAGCAACTGGCCTCTCACTTTCGGCGGAGGGACCAAGGTGGAGATCAAAC

##### M2-4LC

GAAATTGTGTTGACACAGTCTCCAGCCACCCTGTCTTTGTCTCCAGGGGAAAGAGCCACCCTCTCCTGCAGGGCCAGTCAGAGTGTTAGCAGCTACTTAGCCTGGTACCAACAGAAGGTTGGCCAGGCTCCCAGGCTCATCATCTTTGATGCATCCAACAGGGCCACTGGCATCCCAGCCAGGTTCAGTGGCAGTGGGTCTGGGACAGACTTCACTCTCACCATCACCAGCCTAGATCCTGAAGATTTTGCAGTTTATTACTGTCAGCAACGTAGCGCCTGGCCTCTCACTTTCGGCGGAGGGACCAAGGTGGAGATCAGAC

##### M2-5LC

GAAATTGTGCTGACACAGTCTCCAGCCACCCTGTCTTTGTCTCCAGGGGAAAGAGCCACCCTCTCCTGCAGGGCCAGTCAGAGCATTAGCAGCGACTTAACCTGGTTCCAACAGAAACCTGGCCAGGCTCCCAGGCTCATCATCTATGATGCATCCAACAGGGCCACTGGCATCCCAGCCAGGTTCAGTGGCAGTGGGTCTGGGACAGACTTCACTCTCACCATCAGTAGCCTCGAGCCTGAAGATTTTGTAGTTTATTACTGTCTGCAACGTAGCGACTGGCCTCTCACTTTCGGCGGAGGGACCAAGGTGGAGATCAAAC

## Supplementary Figure legends

**Figure S1.**
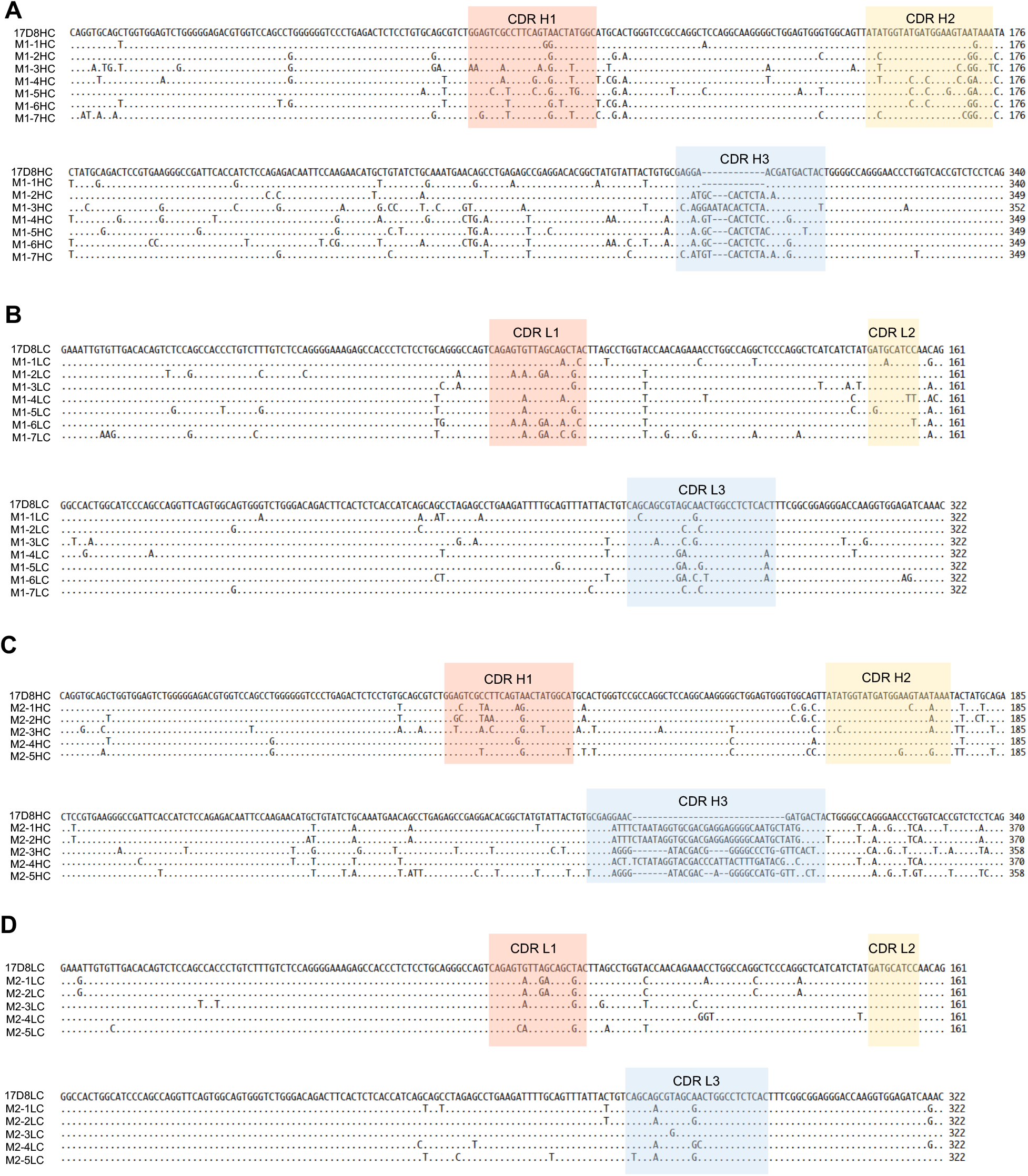
DNA Sequence alignments of the Ig variable regions of new anti-PD1 antibodies with the 17D8HC. **A**. Alignment of M1-1 to M1-7HCs with the 17D8HC. The sequence of the 17D8HC serves as the template for alignment; only SHMs in new anti-PD1 antibodies are shown below, “.” Indicates identity, “−“ indicates gap in alignment. The CDRs are shaded with color. Sections B-D are organized in the same way as this section. **B**. Alignment of M1-1 to M1-7LCs with the 17D8LC. **C**. Alignment of M2-1 to M2-5HCs with the 17D8HC. **D**. Alignment of M2-1 to M2-5LCs with the 17D8LC.

**Figure S2.**
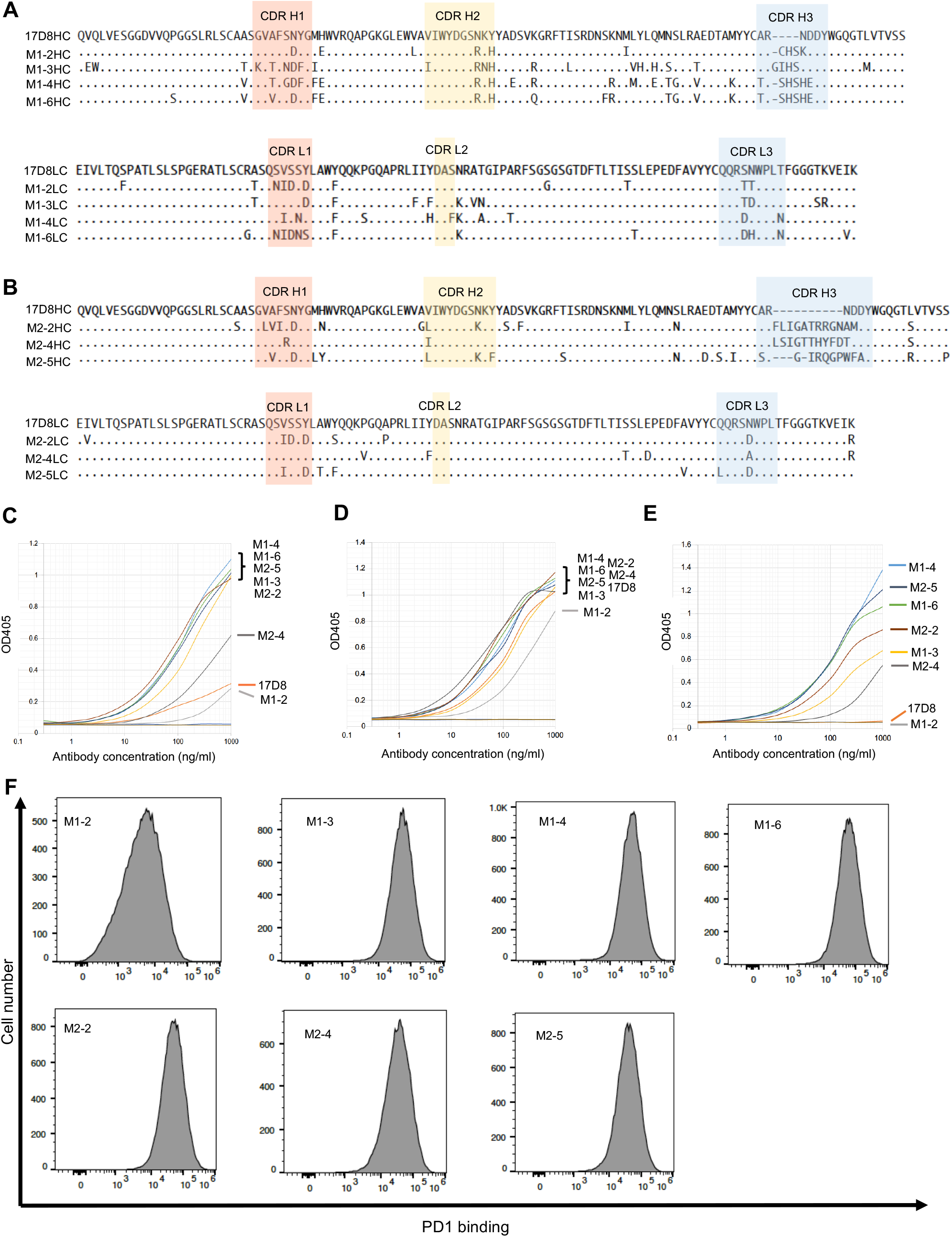
Analysis of PD1-binding activities of new anti-PD1 antibodies. This figure supplements Figure 3 by showing the analysis of additional antibodies isolated from Mouse Models 1 and 2. Sections A-C in this figure correspond to A-C in Figure 3, Section D-F in this figure relate to E-G in Figure 3.

**Figure S3.**
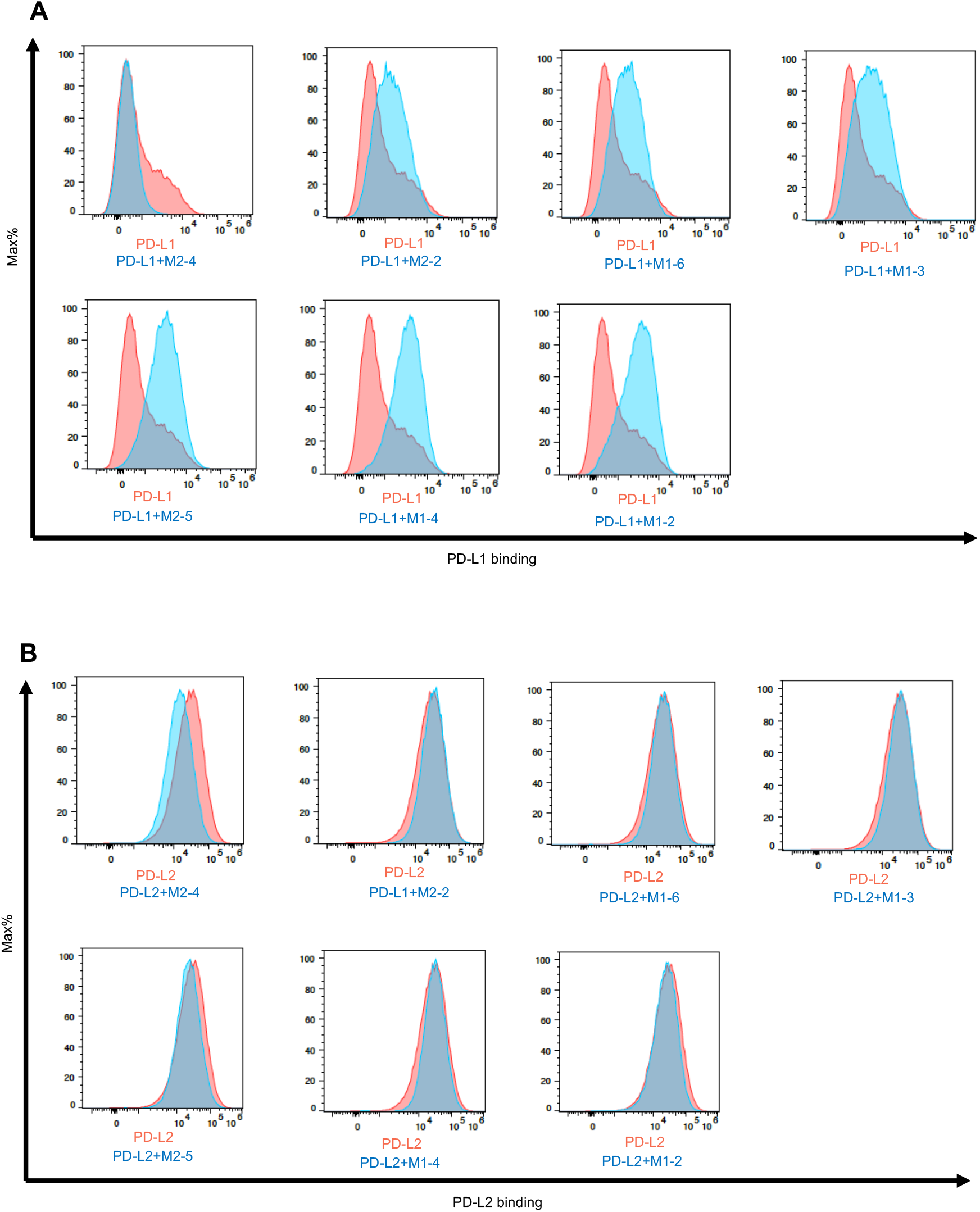
The effects of new anti-PD1 antibodies on PD1/ligand interaction. This figure supplements Figure 4 by presenting the characterization of additional antibodies isolated from Mouse Model 1 and 2. Sections A and B in this figure relate to A and B in Figure 4.

